# Geometric analysis of airway trees shows that lung anatomy evolved to enable explosive ventilation and prevent barotrauma in cetaceans

**DOI:** 10.1101/2024.10.16.618729

**Authors:** Robert L. Cieri, Merryn H. Tawhai, Marina Piscitelli-Doshkov, A. Wayne Vogl, Robert E. Shadwick

**Affiliations:** Department of Zoology, University of British Columbia, Vancouver, BC V6T 1Z4, Canada; The Auckland Bioengineering Institute, Auckland, New Zealand; North Carolina Aquarium at Jannette’s Pier, Nags Head NC, United States

**Keywords:** whales, functional morphology, ventilation, fluid dynamics, lungs, respiratory

## Abstract

Two new biomechanical challenges faced cetacean lungs compared to their terrestrial ancestors. First, hydrostatic pressures encountered during deep dives are sufficient to cause nearly full lung collapse, risking substantial barotrauma during surfacing if air is trapped in the fragile smaller airways. Second, rapid ventilation in large cetaceans requires correspondingly high ventilatory flow rates. In order to investigate how airway geometry evolved in response to these challenges, we characterized airway geometry from 12 species of cetaceans that vary in common dive depth and ventilatory behavior and a domestic pig using computed tomography. After segmenting the major airways, we generated centerline networks models for the larger airways and computed geometric parameters for each tree including mean branching angle, percent volume fraction, and Strahler branching, diameter, and length ratios. When airway geometry was regressed against ventilatory and diving parameters with phylogenetic least squares, neither average branching angle, percent volume fraction, Strahler length ratio or Strahler branching ratio significantly varied with common ventilatory mode or common diving depth. Higher Strahler diameter ratios were associated with slower ventilation and deeper diving depth, suggesting that cetacean lungs have responded to biomechanical pressures primarily with changes in airway diameter. High Strahler diameter ratios lungs in deeper diving species may help to facilitate more complete collapse of the delicate terminal airways by providing for a greater incompressible volume for air storage at depth. On the other hand, lungs with low Strahler diameter ratios would be better for fast ventilation because the gradual decrease in diameter moving distally should keep peripheral flow resistance low, maximizing ventilatory flow rates.

## Introduction

The marine lifestyle of cetaceans imposes their lungs to at least two new biomechanical challenges. First, Cetaceans spend most of their life in apnea beneath the surface of the water and exchange air with the atmosphere during relatively short surface intervals. This is taken to an extreme in dolphins, which use ‘explosive ventilation,’ exchanging up to 90% of total lung capacity in well under a second. Second, diving to depth after inhalation subjects the body and lung to considerable hydrostatic pressure, causing the lung gas to be compressed and expanded in every dive cycle. Diving to 100m should cause an 83% reduction in lung volume, while diving to 500m or more should reduce lung volume by more than 90%. If compressed air is stored in the delicate acinae, it could lead to barotrauma upon expansion. Thus, it is thought that the larger, less compressible airways in deep diving cetaceans act as a safety volume reservoir during diving, storing acinar air so that the acini can empty and collapse fully.

The airways of cetaceans, like all artiodactyls, display monopodial branching, where parent airways generally give rise to a major child branch at an acute angle and a minor child branch at a greater angle. In order to compare airway trees from different species, branches are organized into hierarchical generations or orders depending on the branching pattern. Branches can be organized into generations where counting starts from a proximal stem branch where each child branch is set to (Gen_Parent_ + 1), or Horsfield and Strahler orders, where the peripheral airways are set to order 1 and work backward. Strahler ordering is appropriate for monopodial trees and sets the order of a parent branch one higher than two child branches of the same order or else the same as the child branch of the highest order (Figure 2). Assigning Strahler orders to the airway trees makes it possible to quantify the rate of decrease in branch length or diameter with increasing order (moving from proximal to distal) in the airway tree, or the rate of increase in the number of branches using the Strahler length ratio, Strahler diameter ratio, or Strahler branching ratio (Figure 3). Ratios are calculated as the antilog of the slope of the log(mean diameter in that order) against order.

**Figure 1.**
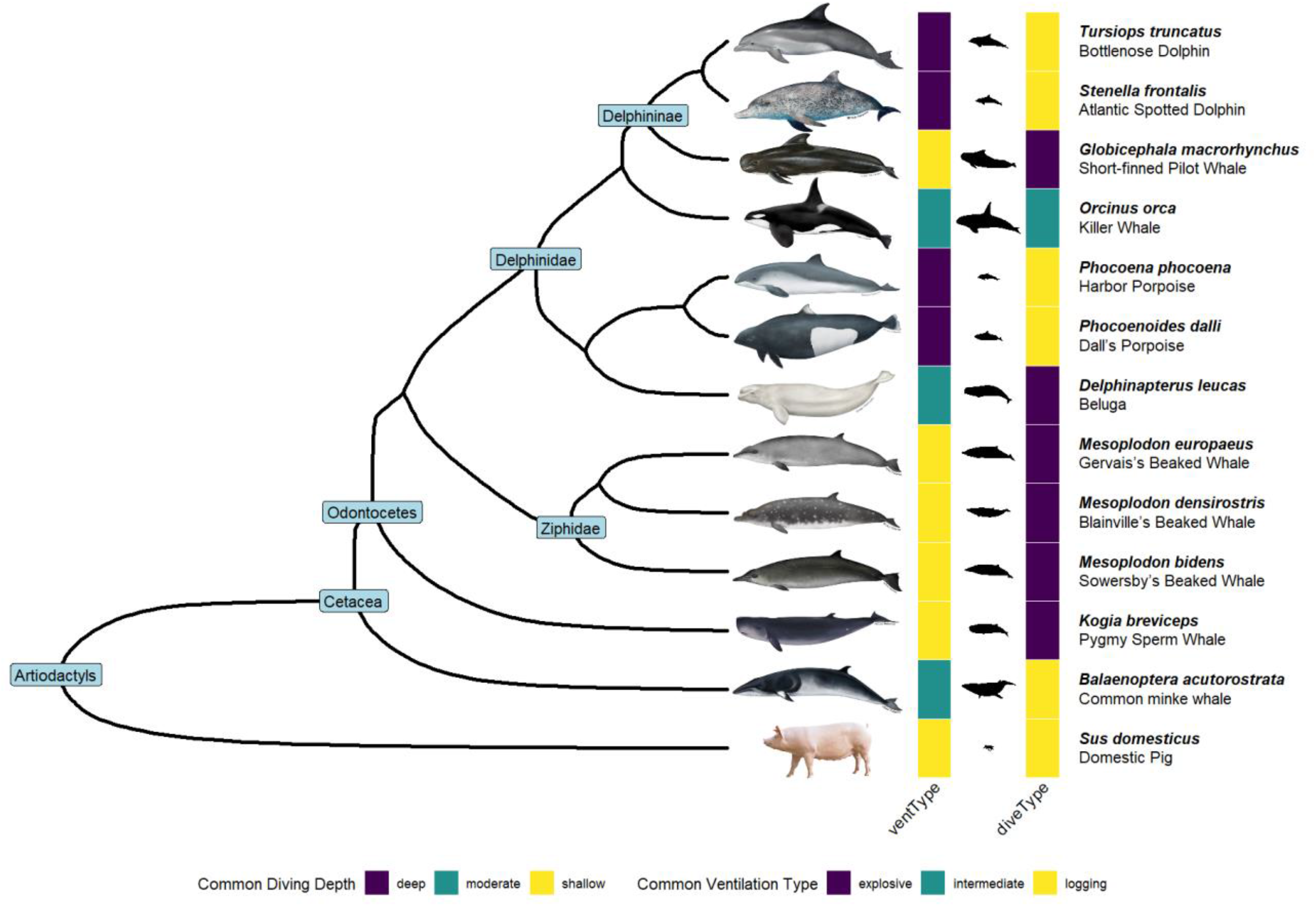
Annotated phylogeny of species included in the study. Phylogenetic relationship of all species included in the study is shown with major clades labeled at node points. Common diving depth and ventilatory types assigned are indicated with colors. Smaller black species silhouettes are scaled to average adult body length. Species illustrations from (NOAA Fisheries 2024).

**Figure 2.**
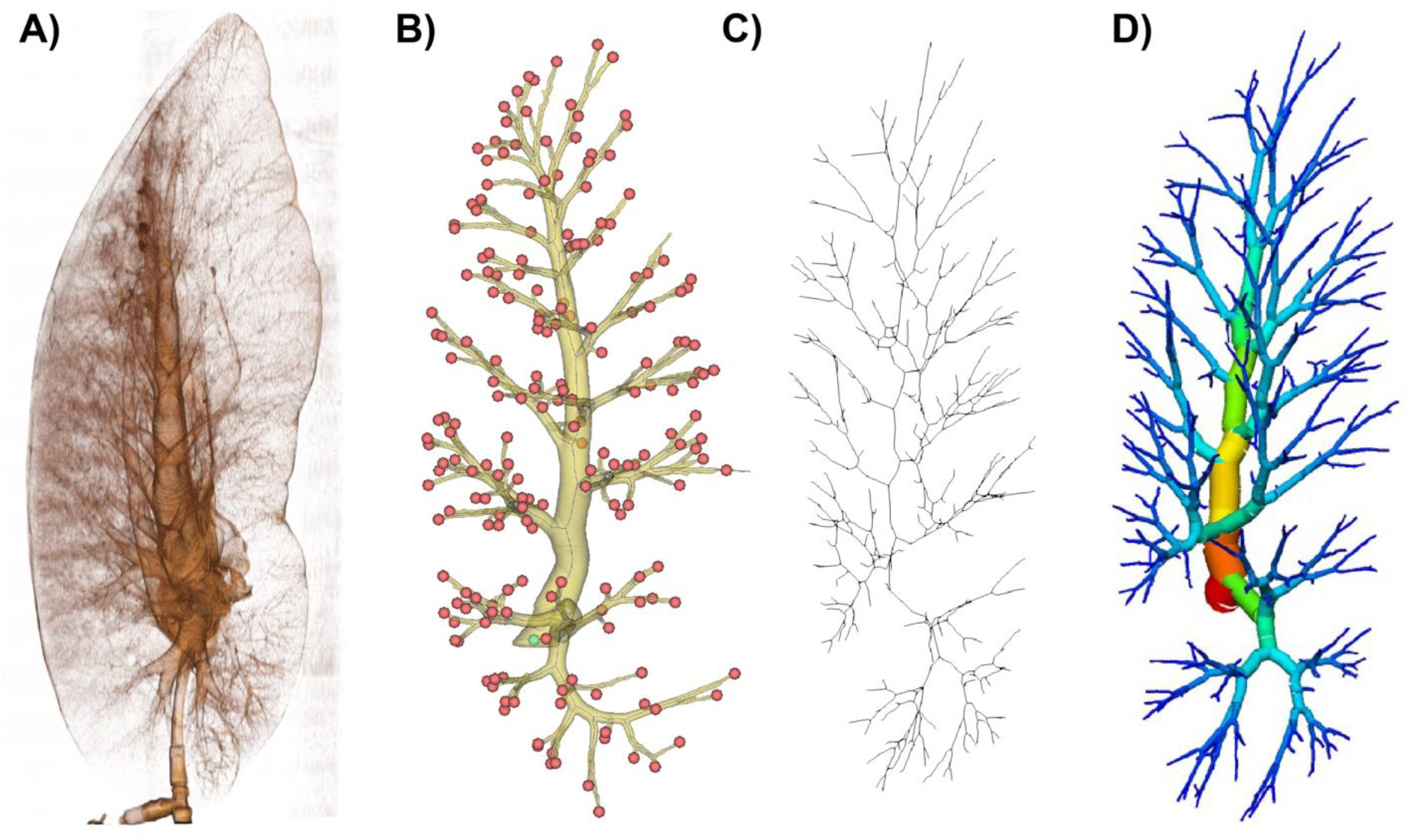
Airway analysis methods pipeline. (A) Volume rendering of lung and major airways for *B. acutorostrata*. (B) Centerline network model (blue lines), centerline endpoints (red circles), and major airway segmentation (yellow). The diameter at network node was computed from the maximum inscribed sphere. (C) Refined one-dimensional airway network tree. (D) Full airway tree model for geometric analysis with topology and segment diameter.

**Figure 3.**
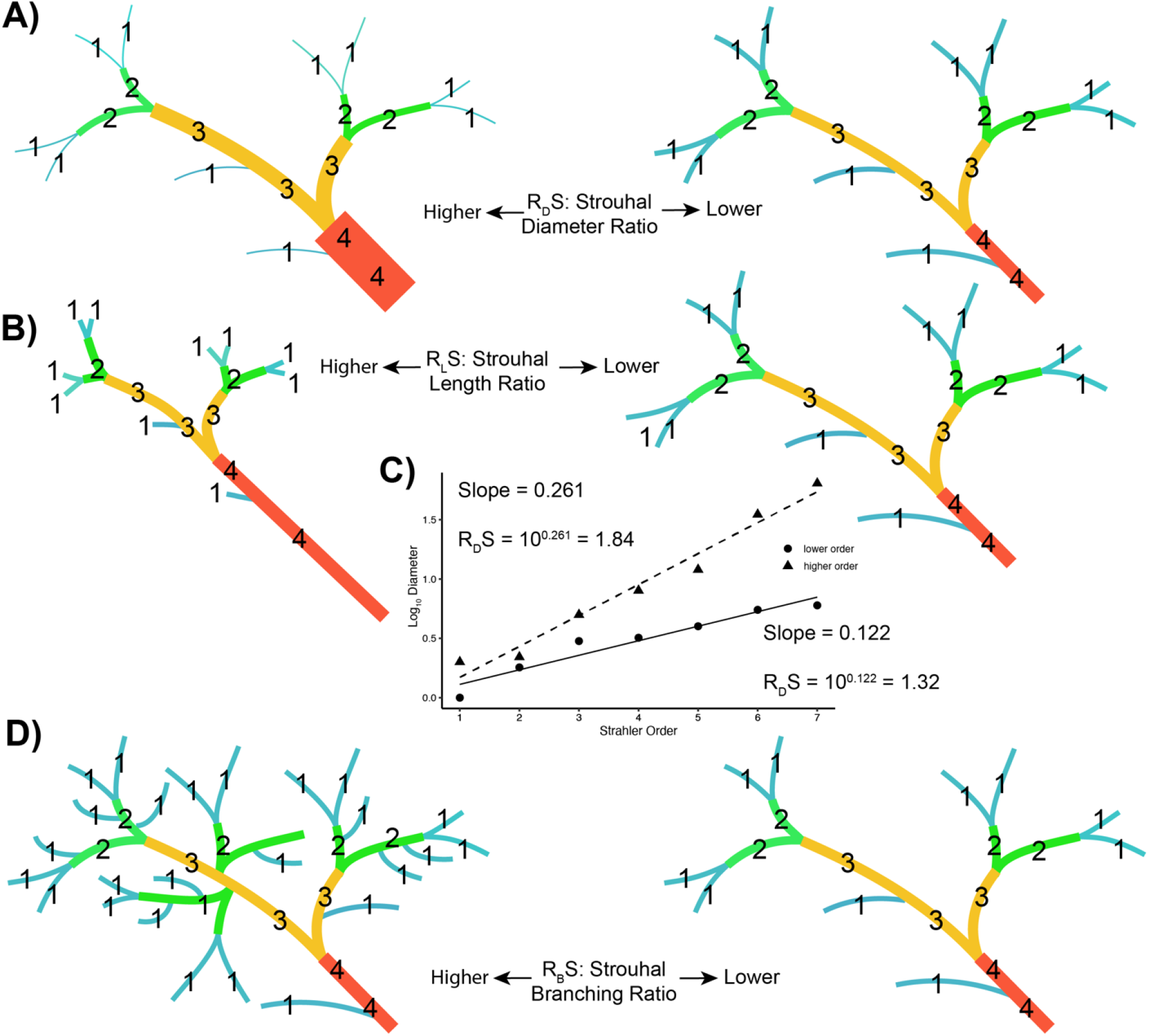
Strahler ordering ratio scheme for airways. Strahler orders start from distal tips and are indicated by numerals on the trees, and color coded. Higher diameter ratios (A) indicate a faster increase in branch diameter with increasing branch order. Higher length ratios (B) correspond to greater increases in branch length with increasing branch order. Plotting of mean airway diameters for hypothetical higher and lower order trees (C) illustrates how ratios are calculated as the antilog of the slopes when the logarithms of values of interest are plotted against branch order. Higher branching ratios (B) correspond to faster decreases in number of branches per order with increasing branch order. Diagram modified from Kilom691 - Own work, CC BY-SA 3.0, https://commons.wikimedia.org/w/index.php?curid=15757078

Thus, we set out to quantify differences in the scaling of airway geometry in the Cetacean pulmonary system to test two contradictory hypotheses: the air storage or barotrauma hypothesis, and the air convection hypothesis. Under the barotrauma hypothesis, relatively wider airways should be found in deeper divers because increased selection pressure against alveolar barotrauma in these species should have provided stronger selection pressure for a greater incompressible volume in the proximal airways. Specifically, we would expect the airway volume fraction, Strahler diameter ratio, and Strahler branching ratio to be larger in deeper-diving species. Under the flow rate hypotheses, relatively larger airways should instead be found in species that use explosive ventilation because the requirements of high flow rates should have led to selection for reduced airway resistance.

## Methods

### Scanning and Data Collection

Lungs from 12 cetacean species plus a domestic pig were acquired through multiple stranding network sources. During necropsy, lungs were excised whole from the carcass either by cutting just below the tracheal carina to remove the left lung intact, or by cutting the tissue around the larynx away to excise both lungs intact including the “goosebeak”. Lungs were then immediately examined while fresh or wrapped in plastic and frozen (−10°C) for later examination. In the case where only the left lung was collected intact, the right lung was examined as standard necropsy protocol and results were reported in the necropsy and histopathology reports along with all other organs.

For computed tomography scanning, the trachea was intubated, and excised lungs were inflated to a set pressure of +30-40 cm H_2_O, representing full total lung capacity (TLC). To minimize movement, the valve was closed, and a high-resolution 3D helical CT scan was performed using a Toshiba Aquilion scanner (Toshiba 3548 Medical Systems Corporation, Tochigi-ken, Japan). The scanner operated with a bed speed of 1 mm/ms and a rotation time of 4 seconds. CT images were captured with a slice thickness of 1.0 mm every 0.5 mm, and the images were reconstructed in the frontal, sagittal, and transverse planes. The resulting in-plane resolution was between 0.5 and 0.3 mm per pixel, with 1 mm spacing between planes, using algorithms optimized for high-resolution bone and soft tissue imaging.

### Behavioral Correlates

For each species, ventilation type was assigned to the most common mode of ventilation (explosive ventilation, logging, or moderate) for that species based on (Piscitelli et al. 2013) and other references. Each species was also assigned to a common diving depth category (deep: dives > 100m are common, shallow: dives < 100m are uncommon, and moderate) based on a literature search. Typical adult body lengths were assigned from (Bisconti, Pellegrino, and Carnevale 2023; Ferguson et al. 2023; Slater, Goldbogen, and Pyenson 2017).

### Anatomical analysis methods

Lung scans were imported in (.*dicom)* format into *ITKsnap* (version 4.2.0, www.ITKsnap.org), an open-source medical imaging software. Major airways (negative air spaces in the lung) were segmented by using the *active contouring* module, which performs level set segmentation (Yushkevich et al. 2006). Briefly, the entire volume was initially segmented using intensity thresholding to exclude voxel densities clearly corresponding to parenchymal tissue, and then between 8-30 seed points were assigned throughout the visible airway tree. The active contouring algorithm was then performed using various sets of control coefficients until as much of the airway tree was segmented as scan resolution permitted. The resulting airway initial segmentations were then exported as surface file (.*stl)*.

Next, initial segmentations and full lung scans were imported into *Dragonfly* (version 2022.2.0.1399, *Object Research Systems*, www.theobjects.com) under a non-commercial use license. The initial surface files were transformed into segmentation layers which were visually inspected and corrected using manual segmentation tools. Refined airway segmentations were re-transformed into surfaces, smoothed using Laplacian smoothing, and exported again as surface files (.*stl)*. The entire lung volume was also segmented to calculate the relative airway volume.

Smoothed airway files and full lung .*dicom* scans were next imported in 3D Slicer, an open-source medical imaging software (Fedorov et al. 2012) (version 5.7.0, www.slicer.org). A centerline network of the segmented airways was calculated using *Extract Centerline* and exported in *VTK* format using the VMTK slicer extension for Slicer3D. *A*irway radius at every network node point was calculated from the maximum inscribed sphere method using *Cross-section analysis* from the VMTK slicer extension for Slicer3D. VTK airway networks were cleaned of duplicate points, formatted into bifurcating airway trees, and assigned Strahler orders using custom python code. Within each Strahler order, branch diameters, counts, and lengths were used to compute branching, diameter, and length ratios.

### Phylogenetic analysis

A time-scaled phylogeny of all species with 13 tip points and 12 internal node points was downloaded from *TimeTree* (Kumar et al. 2017) in Newick format and visualized with the *ggTree* R package (Yu et al. 2017). The tree was annotated with cetacean species images from (NOAA Fisheries 2024) and silhouettes from *Phylopic* (www.phylopic.org) using R packages *ggImage* (Yu 2017) and *rphylpic* (Gearty, Jones, and Chamberlain 2018). Diving depth (deep < moderate < shallow) and ventilatory mode (explosive < moderate < logging) were coded as ordered factors. Phylogenetic least-squared regressions were performed with the function *caper::pgls* (Orme et al. 2011) to determine the influence of diving depth and ventilatory type in a phylogenetic context. Maximum likelihood ancestral state reconstructions along the phylogeny were performed with the function *phytools::fastAnc*. All data were analyzed using R Statistical Software (v4.4.1) (R Core Team 2021) and the *tidyverse* (Wickham 2016), and exported with the R package *flextable* (Gohel and Skintzos 2017) on R Studio 2024.04.2 (www.posit.co).

Colorblind-friendly colors schemes were generated with R packages *viridis* (Garnier 2015) and *scico* (Pedersen and Crameri 2018).

## Results

The percentage of each lung volume consisting of airways, and parenchyma is given in Table 3. Basic geometric parameters calculated from each reconstructed three-dimensional airway trees such as mean branching angle, as well as shape ratios calculated from Strahler orders are also listed in Table 3.

The average branching angle in the airway trees was found to decrease with diving depth (p = 0.049) but increase with faster ventilatory mode (p = 0.009) (Table 1). The volume fraction consisting of airway was not significantly related to any parameters. In terms of Strahler ratios, diameter ratio increased with slower ventilatory mode (p = 0.041), and dive depth (p = 0.029), and length ratio decreased with adult body length (p = 0.024) (Table 2).

**Table 1.** Metadata for individuals included in the study and airway tree characteristics. Percentage of airway volume (airway lung volume / total lung volume ^*^100). Common ventilation type: Common dive depth: shallow (< 100m), deep (> 100m). Airway geometry ratios based on Strahler ordering: branching ratio (R_B_S), diameter ratio (R_D_S),, length ratio (R_L_S). See methods for details.

| species name | adult<br>body length<br>(m) | body mass<br>(kg) | size class | % airway<br>volume | common<br>ventilation type | common<br>dive depth | average<br>branching<br>angle | $R_B S$ | $R_L S$ | $R_D S$ |
| --- | --- | --- | --- | --- | --- | --- | --- | --- | --- | --- |
| Balaenoptera acutorostrata | 6.76 | 902.4 | subadult | 2.36 | intermediate | shallow | 52.994 | 2.56 | 1.22 | 1.492 |
| Delphinapterus leucas | 5.5 | 518.5 | adult | 0.95 | intermediate | deep | 48.474 | 3.446 | 1.165 | 2.189 |
| Globicephala macrorhynchus | 7.2 | 334 | subadult | 1.02 | logging | deep | 50.092 | 2.844 | 1.15 | 2.474 |
| Kogia breviceps | 3.8 | 167 | subadult | 1.48 | logging | deep | 54.955 | 3.575 | 1.379 | 1.433 |
| Mesoplodon bidens | 5.5 | 1014 | adult | 3.09 | logging | deep | 54.821 | 2.986 | 1.535 | 1.518 |
| Mesoplodon densirostris | 4.73 | 340 | subadult | 3.97 | logging | deep | 57.801 | 3.444 | 1.696 | 1.532 |
| Mesoplodon europaeus | 6.71 | 6169 | subadult | 3.76 | logging | deep | 54.227 | 2.814 | 1.487 | 1.577 |
| Orcinus orca | 9.8 | 158 | neonate | 1.81 | intermediate | moderate | 53.027 | 3.036 | 1.287 | 1.775 |
| Phocoenoides dalli | 2.39 | 110.53 | adult | 2.21 | explosive | shallow | 56.662 | 3.434 | 1.633 | 1.502 |
| Phocoena phocoena | 1.86 | 28.8 | subadult | 1.20 | explosive | shallow | 57.17 | 2.653 | 1.602 | 1.893 |
| Stenella frontalis | 2.3 | 241 | adult | 2.48 | explosive | shallow | 57.576 | 2.928 | 1.622 | 1.739 |
| Tursiops truncatus | 4 | 160 | subadult | 1.54 | explosive | shallow | 53.752 | 2.984 | 1.214 | 1.739 |
| Sus domesticus | 1.5 | 36.5 | subadult | 1.09 | logging | shallow | 52.526 | 3.249 | 1.076 | 2.519 |

**Table 2.**
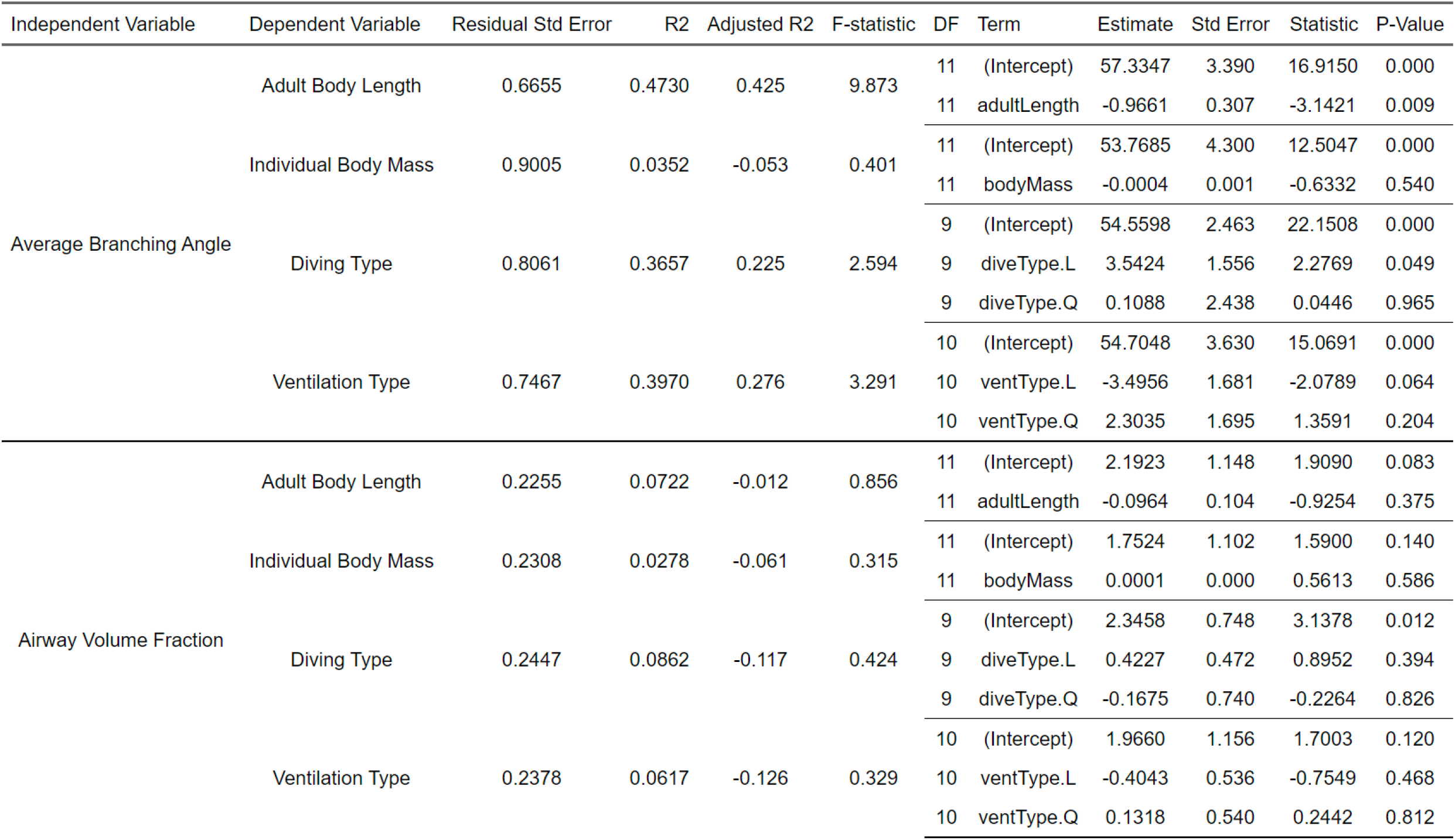
Regression results for branching angle and airway volume fraction. Results of phylogenetic generalized least squared analysis for airway tree geometry parameters regressed against adult body length, body mass of the individual included in the study, common diving depth, and ventilation type. See methods for details.

**Table 3.**
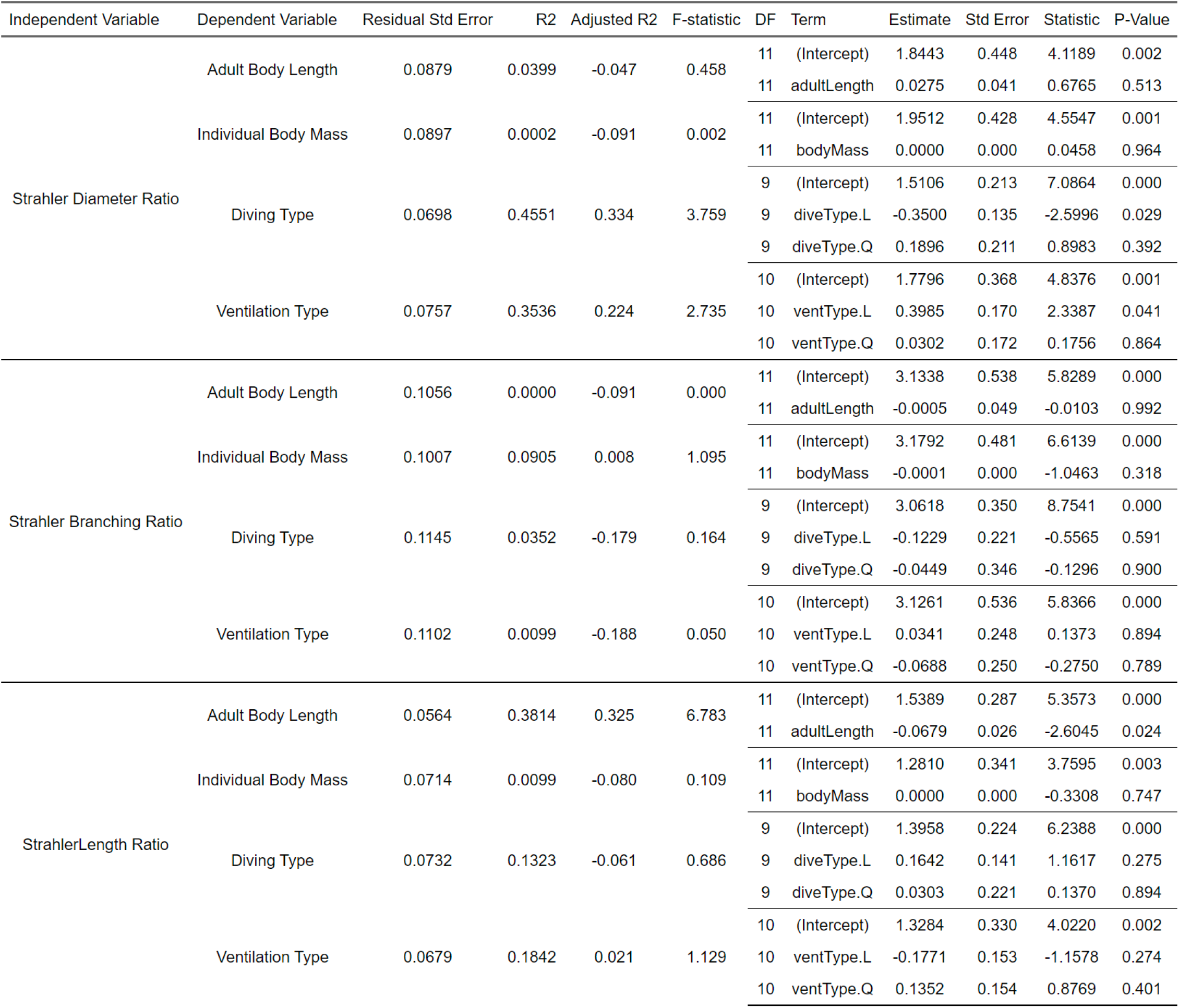
Regression results for branching angle and airway volume fraction. Results of phylogenetic generalized least squared analysis for airway tree Strahler ordering ratios regressed against adult body length, body mass of the individual included in the study, common diving depth, and ventilation type. See methods for details.

## Discussion

We set out to investigate how novel biomechanical challenges associated with evolution towards a marine lifestyle have shaped the pulmonary airway tree in cetaceans. Specifically, we sought to test how the scaling of airway geometry may be influenced by biomechanical pressures associated with explosive ventilation or lung collapse at depth.

The most striking finding of this study is that higher Strahler diameter ratios are associated with deeper diving and logging ventilation in cetaceans. This finding provides strong support for the barotrauma hypothesis because a rapid increase in airway diameter towards the more proximal airways should lead to a greater amount of relatively less-compressible airway volume, promoting more complete collapses of the delicate, smaller airways, preventing distal barotrauma during deep diving. Among delphinids, evolution towards higher diameter ratios are associated with the two deeper-diving and slower ventilating species (Figure 5a), *G. breviceps* and *D. leucas*, providing further support for this association.

**Figure 4.**
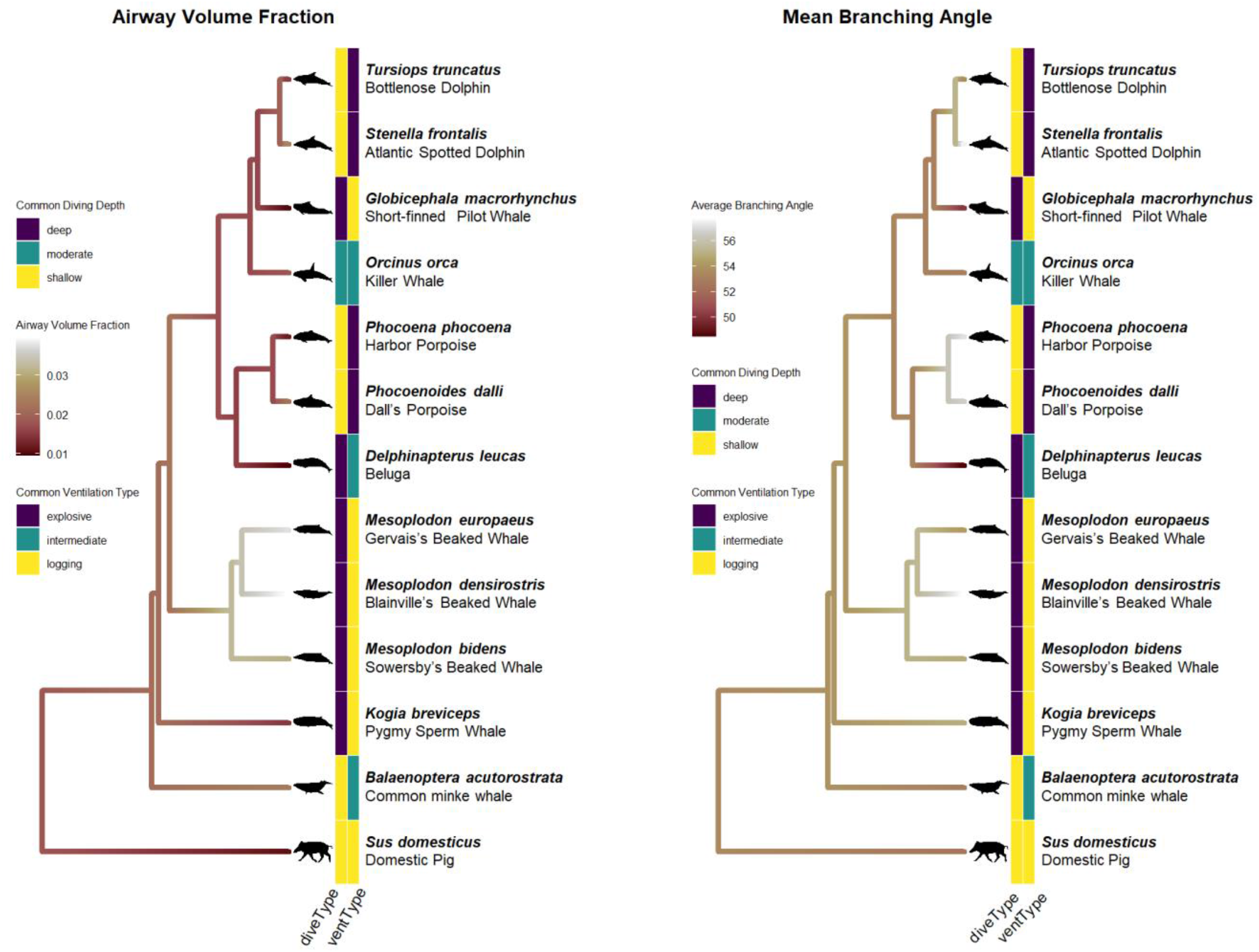
Phylogenetic change in airway volume fraction & mean branching angle. Maximum likelihood ancestral state reconstruction of airway volume fraction (left) and mean airway branching angle (right) plotted on a phylogeny of all species. Crown character states for common diving depth (diveType) and common ventilation type (ventType) are indicated with heatmaps. See methods for details.

**Figure 5.**
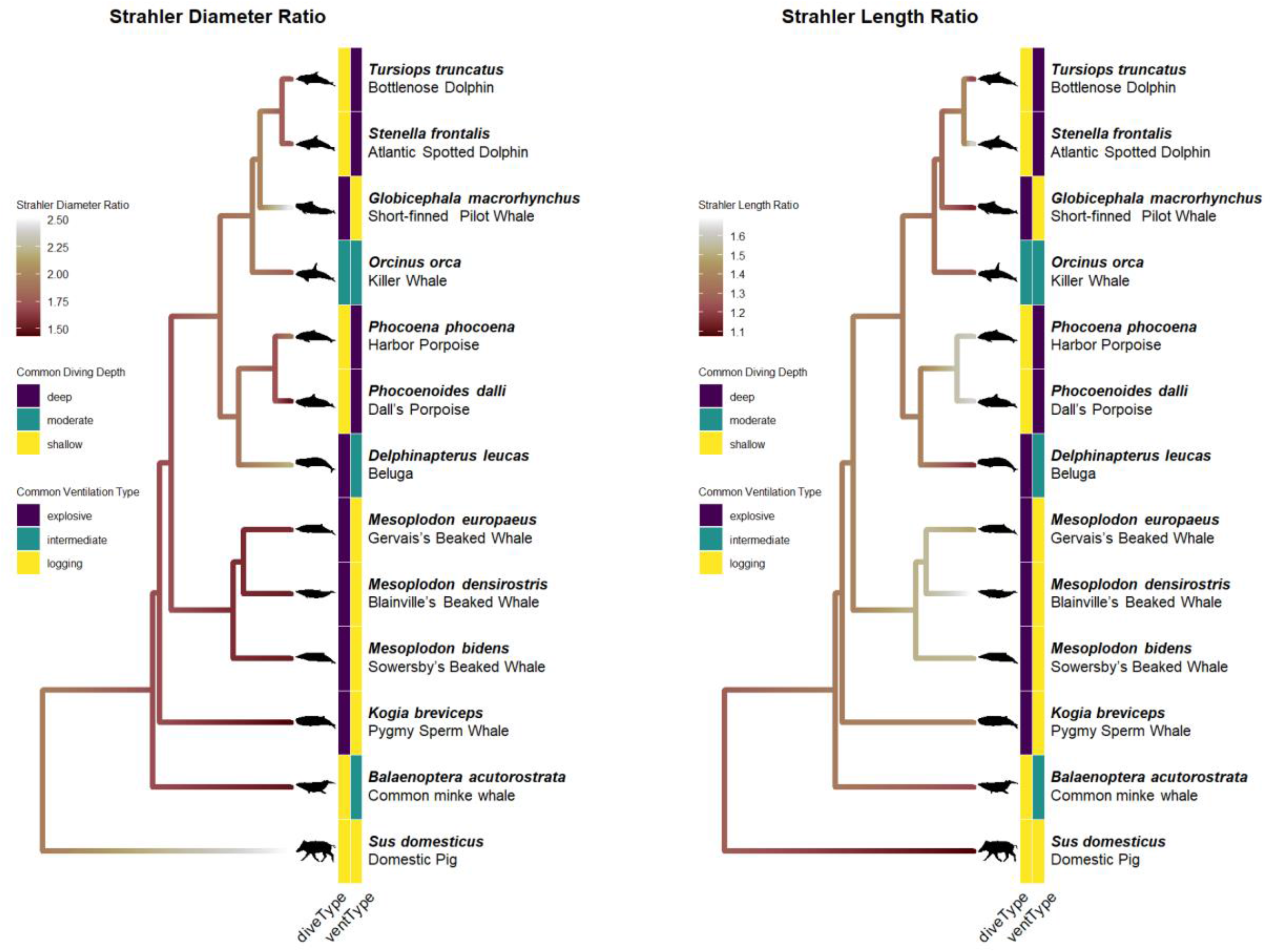
Phylogenetic change in Strahler Diameter Ratio & Strahler Length Ratio. Maximum likelihood ancestral state reconstruction of airway Strahler diameter ratios (left) and Strahler length ratios (right) plotted on a phylogeny of all species. Crown character states for common diving depth (diveType) and common ventilation type (ventType) are indicated with heatmaps. See methods for details.

The association between higher diameter ratios and slow, logging ventilation also provides support for the air convection requirement, because lungs with higher diameter ratios have a greater proximal-to-distal resistance gradient. Most of the resistance to flow in the pulmonary tree is due to the smaller airways (Tawhai and Lin 2011), so faster ventilatory cycles should provide selection to lower diameter ratios.

Interestingly, it seems that beaked whales (*Ziphidae*) have evolved different pulmonary adaptations associated with deep diving than either kogiids or pilot whales, although this conclusion is tentative because each of the latter groups is only represented by one species in this study (*Kogia breviceps*, and *Macrorhynchus globicephala*, respectively). Beaked whales are the deepest-diving whales, spending much of their time below 500m in depth (Pitman 2018), and have evolved a much higher airway volume fraction than other cetaceans (Figure 4a), suggesting that this trait may only be selected for in extreme divers. In contrast, *K. breviceps* has a noticeably high branching ratio, so perhaps sufficient air storage volume is accomplished in kogiids with more moderately sized airways instead of a few major airways. If so, this phenotype – which seems to be shared with another transitional species, *D. leucas*, may represent a trade-off between preventing barotrauma and limiting peripheral resistance to enable faster ventilatory flow rates.

Although no significant relationship between mean branching angle and either diving depth of ventilatory type was detected in cetaceans overall (Table 1), higher branching angles seem to be associated with explosive ventilation and shallow diving among delphinids (Fig 4b), although these are also the smallest species. Higher branching angles may facilitate complete lung emptying during explosive ventilation because a greater proportion of lung gas would flow into the major bronchi through proximal openings. Similarly, higher branching angles were also found in beaked whales, which could help to facilitate complete movement of lung gasses from smaller to larger airways during diving.

Although only one baleen whale is represented, it seems that airway geometry is relatively intermediate in these animals, although *B. acutorostrata* has a notably low branching ratio (Figure 6a). If this is true for other, larger, rorqual whales, it may suggest that the larger terminal airway structures found in rorquals (Engel 1966) may be connected more directly to the major bronchi than in other cetaceans. It is unknown how airway geometry will change with the extreme body size seen in larger rorquals.

**Figure 6.**
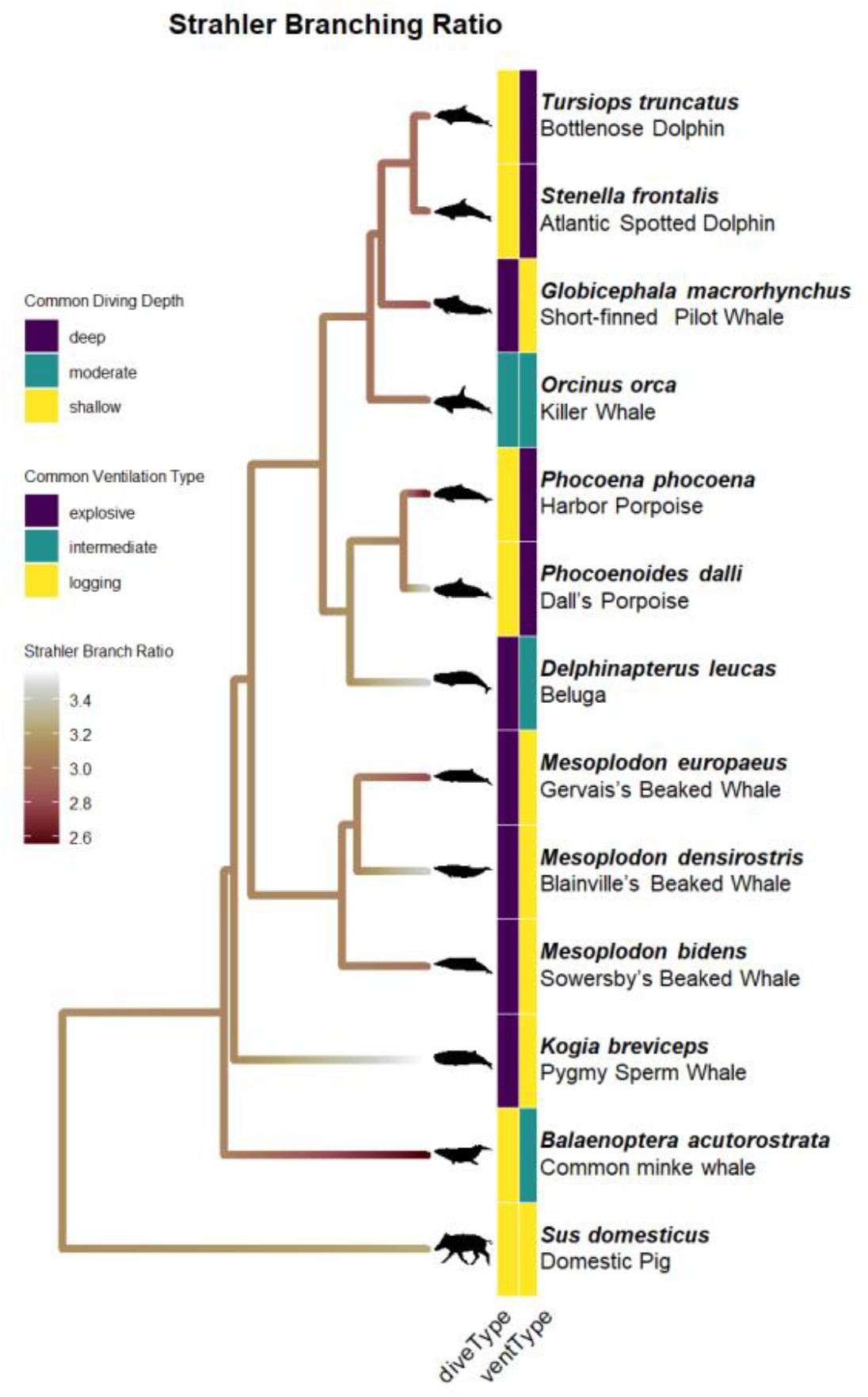
Phylogenetic change in Strahler Branching Ratio. Maximum likelihood ancestral state reconstruction of airway Strahler branching ratios plotted on a phylogeny of all species. Crown character states for common diving depth (diveType) and common ventilation type (ventType) are indicated with heatmaps. See methods for details.

This study also provides insight into how the lung has evolved in cetaceans generally. Between outgroup artiodactyls and stem cetaceans, SDR increased substantially (Figure 5a), indicating that cetaceans evolved may have evolved this trait with the initial radiation into the water, but tis change may be the result of larger body sizes in cetaceans. Some cetaceans have somewhat larger terminal airway volumes than terrestrial mammals (Piscitelli et al. 2013), but the major airways are presumably much more relatively large, so the SDR must be greater in cetaceans unless they have many more airway generations. Future work examining the geometry of the cetacean airway tree down to the acinar level, or investigation of airway tree geometry in other large mammals such as elephants, would be a good test of this hypothesis.

## Acknowledgements

This work was supported by Natural Sciences and Engineering Research Council of Canada Discovery Grant RGPIN-10 2019-04235. This material is based upon work supported by the NSF Postdoctoral Research Fellowships in Biology Program under Grant No. (2010666). Any opinions, findings, and conclusions or recommendations expressed in this material are those of the author(s) and do not necessarily reflect the views of the National Science Foundation.

